# Prolonged airway explant culture enables study of health, disease, and viral pathogenesis

**DOI:** 10.1101/2024.02.03.578756

**Authors:** Rhianna E Lee-Ferris, Kenichi Okuda, Jacob R Galiger, Stephen A Schworer, Troy D Rogers, Hong Dang, Rodney Gilmore, Caitlin Edwards, Satoko Nakano, Anne M. Cawley, Raymond J Pickles, Samuel C Gallant, Elisa Crisci, Lauraine Rivier, James S Hagood, Wanda K O’Neal, Ralph S Baric, Barbara R Grubb, Richard C Boucher, Scott H Randell

## Abstract

In vitro models play a major role in studying airway physiology and disease. However, the native lung’s complex tissue architecture and non-epithelial cell lineages are not preserved in these models. Ex vivo tissue models could overcome in vitro limitations, but methods for long-term maintenance of ex vivo tissue has not been established. We describe methods to culture human large airway explants, small airway explants, and precision-cut lung slices for at least 14 days. Human airway explants recapitulate genotype-specific electrophysiology, characteristic epithelial, endothelial, stromal and immune cell populations, and model viral infection after 14 days in culture. These methods also maintain mouse, rabbit, and pig tracheal explants. Notably, intact airway tissue can be cryopreserved, thawed, and used to generate explants with recovery of function 14 days post-thaw. These studies highlight the broad applications of airway tissue explants and their use as translational intermediates between in vitro and in vivo studies.

## Introduction

Airway epithelia constitute an important barrier between the body and the environment, serving as the first point of contact with inhaled chemicals, particles, and pathogens. Airway barrier function is crucial for health, and barrier failure often leads to disease. Air-liquid interface (ALI) cultures have long served to model airway epithelial barrier function and physiology^1,2^. In this system, airway epithelial cells form a pseudostratified epithelium containing the major cell types of in vivo airways, including basal, secretory, and ciliated cells. However, in vivo stromal architecture, submucosal glands, and epithelial interactions with tissue-resident immune cells are not preserved. Animal models offer another strategy to study the respiratory epithelium. However, translating findings from animal models to humans is not always straightforward.

Airway explants contain diverse cell types and spatial information characteristic of the in vivo tissue and, thus, may serve as an intermediate between ALI and in vivo models. Despite these advantages, explant culture methods typically require fresh tissue obtained within a few hours of resection^3^. Methods to recover and culture human tissues after longer cold ischemic times have not been developed. Further, explants have been historically limited by short-term viability (∼4 days)^4–8^ and submerged culture methods, which do not mimic the in vivo air–tissue interface.

Here, we developed methods to culture explants from three major lung regions, including bronchi (large airway explants; LAE), bronchioles (small airway explants; SAE), and alveolar parenchyma (precision cut lung slices; PCLS). Maintaining explants on a Gelfoam® sponge enabled an air-liquid interface, circumventing the need for submerged culture. Even after long cold ischemic times (up to 40h), cultured airway explants remained viable and retained in vivo-like characteristics for 14 days. We assessed explant cellular composition, electrophysiology, cryopreservation and demonstrated their utility to study viral infection.

## Results

*Gelfoam® culture maintains in vivo-like tissue architecture and characteristic cell types.* The epithelium of human LAE and SAE was examined immediately after dissection and after 14d of culture on Gelfoam® at an air-liquid interface (see methods) (Fig. 1A-F). Much like the freshly explanted tissue (Fig. 1A-B), the d14 LAE and SAE epithelium was pseudostratified and well-ciliated, with submucosal glands containing AB-PAS positive cells and secretions visible in LAE (Fig. 1C-F). By immunofluorescence (IF), abundant acetylated α-tubulin+ ciliated cells and sparse mucin 5AC (MUC5AC)+ goblet cells were observed in the surface epithelium, along with robust mucin 5B (MUC5B) staining in the submucosal glands of d14 LAE (Fig. 1G). Bundles of actin alpha 2, smooth muscle (ACTA2)+ smooth muscle cells, platelet and endothelial cell adhesion molecule 1 (PECAM1)+ endothelial cells, and protein tyrosine phosphatase receptor type C (PTPRC)+ leukocytes were present below the epithelium (Fig. 1H). Macrophages, positive for cluster of differentiation (CD) 68 (CD68), T cells (CD3+), and B cells (CD20+) were also present in d14 LAE tissues (Fig. I-J). SAE at d14 also expressed α-tubulin, MUC5B, MUC5AC, ACTA2, PECAM1, and PTPRC (Fig. 1K-L) and retained viable alveolar tissue on the Gelfoam®-facing side of the explant, as indicated by IF for the alveolar type 2 (AT2) cell markers lysosomal associated membrane protein 3 (LAMP3) and pro-surfactant protein B (pro-SPB) and the alveolar type 1 (AT1) cell marker, advanced glycosylation end-product specific receptor (AGER) (Fig. 1M).

**Figure 1:**
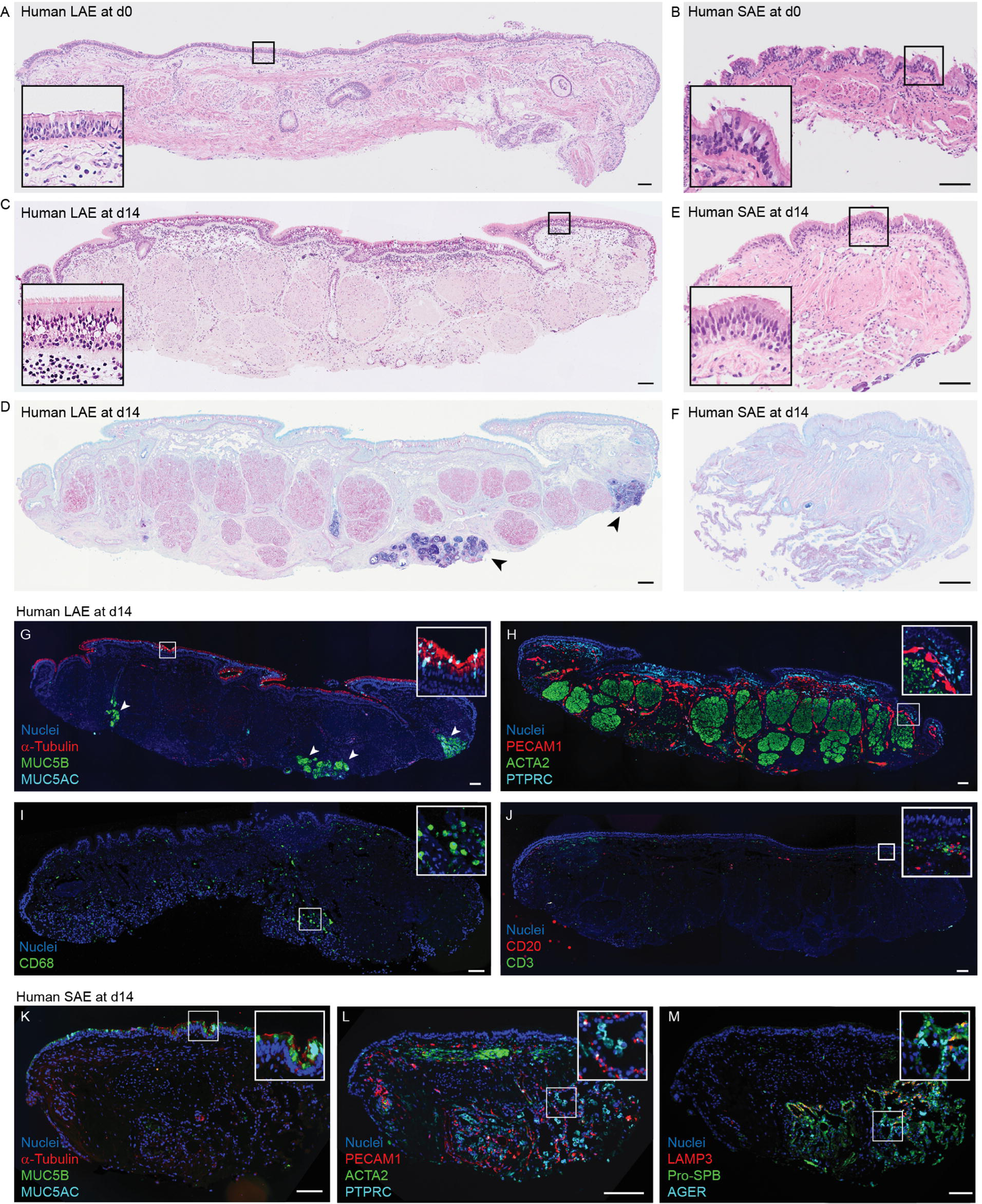
Human LAE and SAE exhibit native tissue architecture and characteristic epithelial, mesenchymal, and immune cell types after 14 days in culture. A-B) H&E staining of human LAE (A) and SAE (B) immediately after dissection (d0). Representative of N = 7 and 5 donors, respectively. C-D) H&E (C) and AB-PAS (D) staining of a human LAE explant after 14 days in culture. Arrows in (D) indicate submucosal glands. Representative of N = 17 donors. E-F) H&E (E) and AB-PAS (F) of a human SAE explant after 14 days in culture. Representative of N = 13 donors. G-J) Immunofluorescence localization of α-tubulin, MUC5B, and MUC5AC (G); PECAM1, ACTA2, and PTPRC (H); CD68 (I); and CD20 and CD3 (J) in d14 LAE explants. Arrows in (G) indicate submucosal glands. Representative of N = 3 donors. K-M) Immunofluorescence localization of α-tubulin, MUC5B, and MUC5AC (K); PECAM1, ACTA2, and PTPRC (L); and LAMP3, Pro-SPB, and AGER (M) in d14 SAE explants. Representative of N = 3 donors. All scale bars = 100 µm.

Maintenance of characteristic airway epithelial cells was confirmed by RNA *in situ* hybridization. Abundant *Forkhead box J1 (FOXJ1)* signal, marking ciliated cells, and *Mucin 5B (MUC5B)* signal marking secretory cells in the surface epithelium of LAE and SAE as well as in the gland ducts and acini in LAE tissues were observed (Extended Data Fig. 1A-B; 1E-F). The secretory cell marker, *Secretoglobin Family 1A Member 1* (*SCGB1A1*), was also present in LAE and SAE tissues, with higher abundance in SAE as previously reported^9^ (Extended Data Fig. 1C; 1G). *Keratin 5* (*KRT5*), a marker of airway basal cells, was present in the surface epithelium and gland ducts of LAE (Extended Data Fig. 1D). We also detected expression of *Surfactant Protein B (SFTPB),* a marker of distal airway secretory cells and AT2 cells, in both the airway surface epithelium and alveolar parenchyma of SAE tissues (Extended Data Fig. 1H). Overall, LAE and SAE appeared to preserve characteristic epithelial, endothelial, and immune cell populations for at least 14 days in culture.

*Human precision cut lung slices cultured on Gelfoam® preserve native tissue architecture and characteristic cell types.* In vitro model systems to study the alveolar epithelium over extended time intervals are less established compared to the upper airways. Precision cut lung slices (PCLS) preserve the alveolar epithelium within the context of its native complex tissue architecture and have been widely adopted to study airway constriction, toxicology, nanotechnology, and viral pathogenesis^10–14^. However, most PCLS studies are limited to short-term culture (≤72h)^12,15,16^. We hypothesized that culturing PCLS on Gelfoam® at an air-liquid interface would extend tissue viability while maintaining in vivo tissue functional properties. H&E staining of d0 and d14 Gelfoam®-cultured PCLS revealed preservation of intact alveolar structures (Fig. 2A-B). In some regions of d14 PCLS, larger patches of cells were seen in the alveolar interstitium (Fig. 2B, arrows), reflecting increased numbers of an unidentified cell type. The viability of PCLS cultured on Gelfoam® and under conventional submerged culture conditions was measured using a WST-8 cell counting kit (Fig. 2C). Submerged PCLS viability began decreasing after 4 days in standard culture conditions, whereas the viability of PCLS on Gelfoam® remained stable for 21 days before decreasing. Collectively, these data indicate that air-liquid interface culture of PCLS on Gelfoam® was superior to conventional submerged culture methods and this method was used for all remaining experiments.

**Figure 2:**
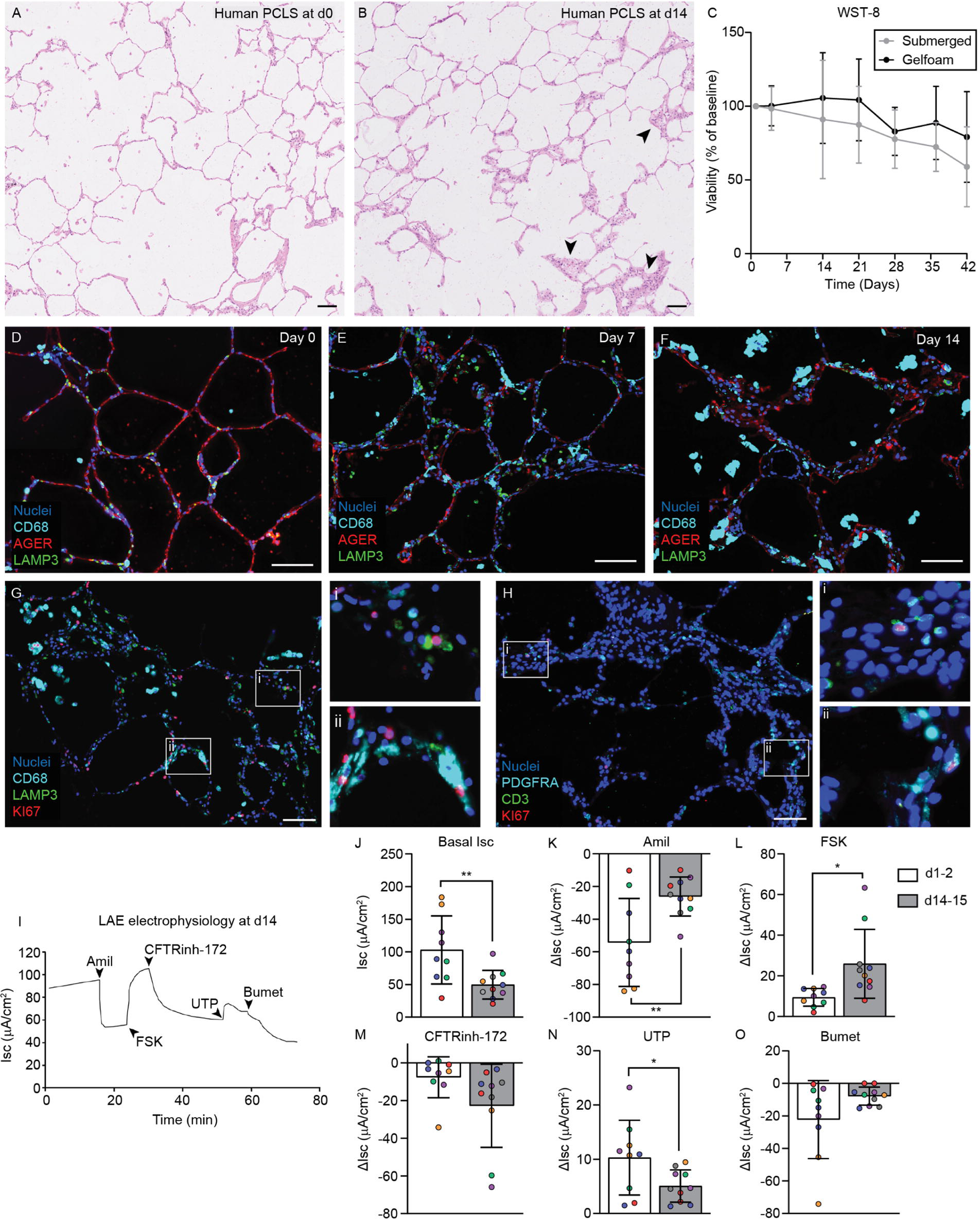
Human PCLS and LAE remain viable for 14 days of culture on Gelfoam. A-B) H&E histology of a d0 (A) and d14 (B) human PCLS. Representative of N = 8 donors. C) WST-8 viability assessment of human PCLS cultured at air-liquid interface on Gelfoam or under traditional submerged methods. N = 3 donors; 3 replicates per donor. Gelfoam-cultured PCLS had significantly greater viability using a linear mixed-effect model with the donor as a random effects factor. D-F) Immunofluorescent localization of CD68, AGER, and LAMP3 in human PCLS at d0 (D), d7 (E), and d14 (F). Representative of N = 3 donors. G-H) Immunofluorescent localization of KI67 with CD68 and LAMP3 (G) and with CD3 and PDGFRA (H) in d14 human PCLS. Representative of N = 3 donors. All scale bars = 100 µm. I-O) Electrophysiology of human LAE explants at d1-2 and d14-15. I) Representative Ussing tracing of a human LAE explant at d14. J) Basal Isc and ΔIsc in response to K) amiloride (Amil), L) forskolin (FSK), M) CFTRinhibitor-172 (CFTRinh-172), N) uridine-5’-triphosphate (UTP), and O) Bumetanide (Bumet). N = 5-6 donors (represented by different colored dots); 1-2 replicates per donor. Unpaired t-test; * = p<0.05, ** = p<0.01.

Next, the cell type composition of PCLS cultured for 0, 7, and 14 days was assessed by IF (Fig. 2D-F). At all three time points, CD68+ macrophages, AGER+ AT1 cells, and LAMP3+ AT2 cells were observed. RNA *in situ* hybridization further confirmed *CD68, AGER,* and *SFTPB* expression in d14 PCLS (Extended Data Fig. 1I-K). We again noted increased cellularity in the alveolar interstitium as well as a marked increase in CD68+ macrophages from d0 to d14 (Fig. 2D-F). Thus, we performed IF localization for the proliferation marker Ki67 in d0 and d14 tissues. Ki67 signal was nearly absent from d0 tissues (Extended Data Fig. 1L-M), but colocalized with a subset of CD68+ macrophages (Fig. 2G), LAMP3+ AT2 cells (Fig. 2G), CD3+ T cells (Fig. 2H), and PDGFRA+ fibroblasts (Fig. 2H) in d14 PCLS.

*LAE and SAE exhibit in vivo-like electrophysiology after 14 days in culture.* Explant viability and function was investigated by assessing electrophysiology in Ussing chambers. There was no measurable electrophysiology immediately after dissection, likely due to the long cold ischemic times of the studied tissues (up to 40h between surgical resection and placement on Gelfoam*®*). However, LAE and SAE electrophysiology recovered after 1-2d or 14-15d in culture (Fig. I-O; Extended Data Fig. 2A-G). When equilibrated in bilateral Krebs Bicarbonate Ringers (KBR) solution, the mean basal short circuit current (Isc) of LAE and SAE were 112.1 and 42.4 µA/cm^2^, respectively at d1-2, and 49.7 and 56.7 µA/cm^2^, respectively at d14-15 (Fig. 2J; Extended Data Fig. 2B). The d14-15 measurements were strikingly similar to measurements from previous studies of freshly excised bronchi and bronchioles which exhibited basal Isc of 51 and 55 µA/cm^2^, respectively^8,17^. By contrast, the two-fold greater basal Isc and amiloride response along with the muted forskolin response at d1-2 likely reflects ongoing recovery from cold ischemia following surgical resection.

In d14-15 explants, robust responses to amiloride (Amil) indicated epithelial sodium channel (ENaC) activity, and responses to forskolin (FSK) and CFTR inhibitor-172 (CFTRinh-172) indicated cystic fibrosis transmembrane regulator (CFTR) activity (Fig. 2K-M; Extended Data Fig. 2C-E), again aligning with previous work^8,17^. Lastly, we compared CF and non-CF LAE after 14d of culture and observed no CFTR activity in CF LAE as expected, indicating that the disease phenotype of reduced CFTR function was preserved by explant culture (Extended Data Fig. 2H-N). Collectively, these data suggest that Gelfoam® culture methods enable culture of viable airway explants which model *in vivo*-like electrophysiology after 14d in culture.

*Single cell RNA sequencing reveals diverse cell lineages preserved in the LAE, SAE, and PCLS models.* To query cell population dynamics in the Gelfoam® model, single-cell RNA-sequencing (scRNA-seq) was performed on LAE, SAE, and PCLS from N=3 donors. After 0, 2, or 14 days in culture, samples were preserved in CryoStor commercial freezing media and processed as one batch per donor before combining and integrating the data (Fig. 3A; Extended Data Fig. 3). Using the quality control metrics defined in the methods, we obtained a total of 85,305 cells combining all three regions and time points. A graph-based clustering approach using Seurat v4 produced 33 distinct clusters (Fig. 3B). We then referenced the Human Lung Cell Atlas dataset^18^ to assign prediction scores for 61 distinct cell signatures. Clusters 3, 6, 22, 23, and 32 contained cells from a single donor only and were not considered representative of the model as a whole (Extended Data Fig. 3F).

**Figure 3:**
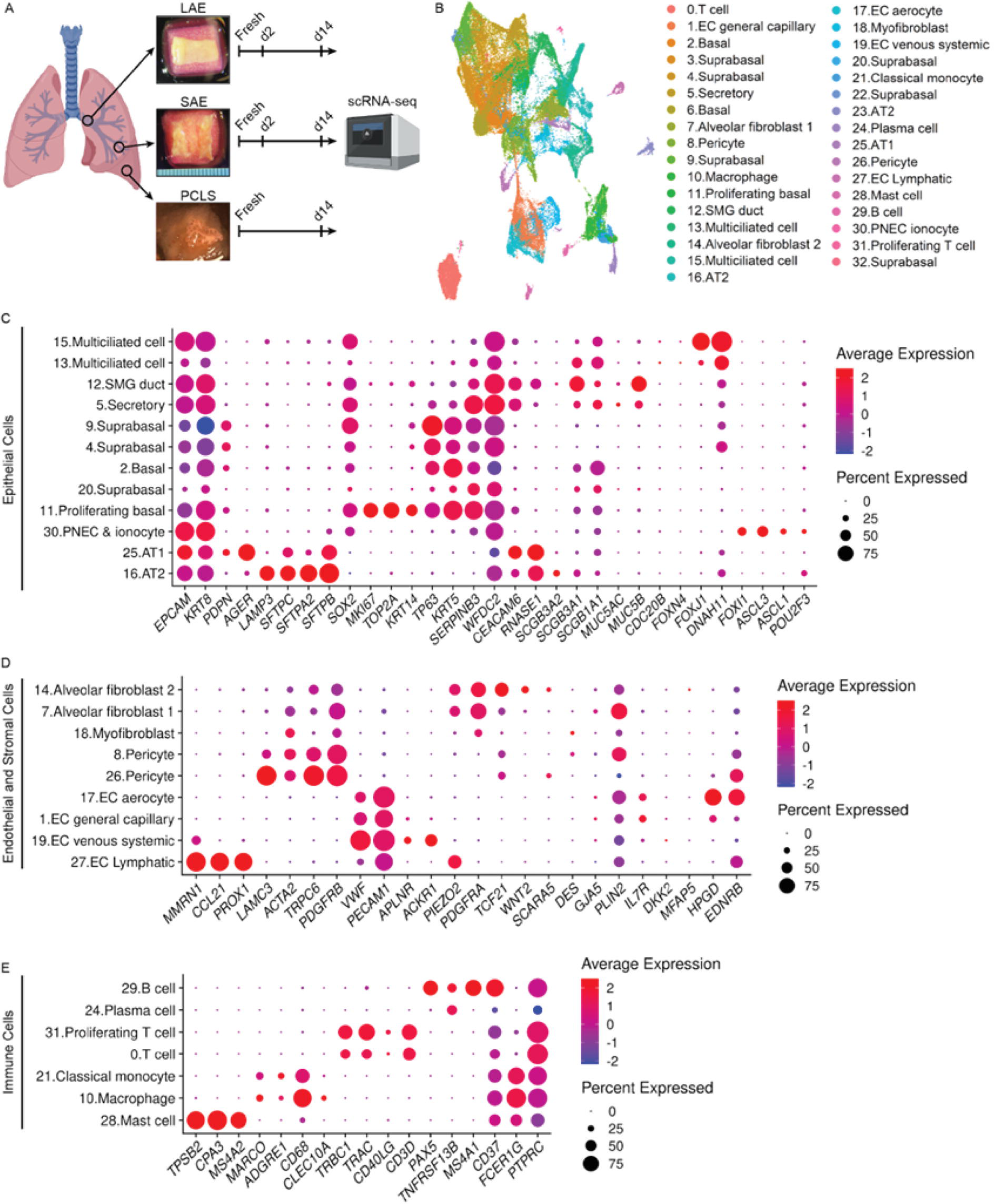
Single cell RNA-sequencing (scRNA-seq) of LAE, SAE, and PCLS models over time. A) Schematic of scRNA-seq on LAE, SAE, and PCLS at d0 (fresh), d2, and d14. B) UMAP indicating 33 identified cell clusters. C-E) Dot plot showing the top differentially expressed genes in epithelial (C), endothelial and stromal (D), and immune cell (E) clusters.

Multiple epithelial cell clusters, including basal cells, suprabasal cells, secretory cells, submucosal gland (SMG) duct cells, multiciliated cells, AT1s, AT2s, pulmonary neuroendocrine cells (PNECs), and ionocytes were identified in our data set based on characteristic marker expression (Fig. 3C). Endothelial cell (EC) clusters included EC capillary, venous systemic ECs, lymphatic ECs, and the recently described EC aerocytes^19^ (Fig. 3D). Stromal cell clusters included fibroblasts, myofibroblasts, pericytes, and alveolar fibroblast 1 and 2 (AF1 and AF2, respectively) (Fig. 3D). Finally, immune cell clusters included macrophages, T cells, B cells, mast cells, plasma cells, and classical monocytes (Fig. 3E).

Our data identified 5 distinct populations of basal or suprabasal cells (Fig. 4A-E). These clusters were generally more abundant in LAE than SAE explants, and virtually absent from PCLS, reflecting the small numbers of conducting airways in PCLS. These clusters expressed the basal cell markers *KRT5* and *KRT15*, as expected (Extended Data Fig. 4A-B). Cluster 2 (basal) decreased over time (Fig. 4A), possibly representing a homeostatic cell state that is not preserved by explant culture methods. By contrast, cluster 11 (proliferating basal) transiently increased at d2 but decreased again by d14 (Fig. 4B) and expressed high levels of the proliferation markers, *MKI67* and *TOP2A* (Extended Data Fig. 4C-D). This pattern was also observed by IF localization of KI67 with abundant signal at d2 and no signal at d15 (Extended Data Fig. 4E-F). Clusters 4 and 9 (suprabasal) increased over time (Fig. 4C-D), and expressed relatively high levels of the hypoxia markers, *EGLN3* and *P4HA1*^20^ (Extended Data Fig. 4G-H). This may indicate an epithelial response to apical hypoxia over time, potentially explained by the visible accumulation of apical mucus (Extended Data Fig. 4I). Cluster 20 (suprabasal) cell abundance was relatively constant in LAE, but transiently increased at d2 in one donor in the SAE model (Fig. 4E).

**Figure 4:**
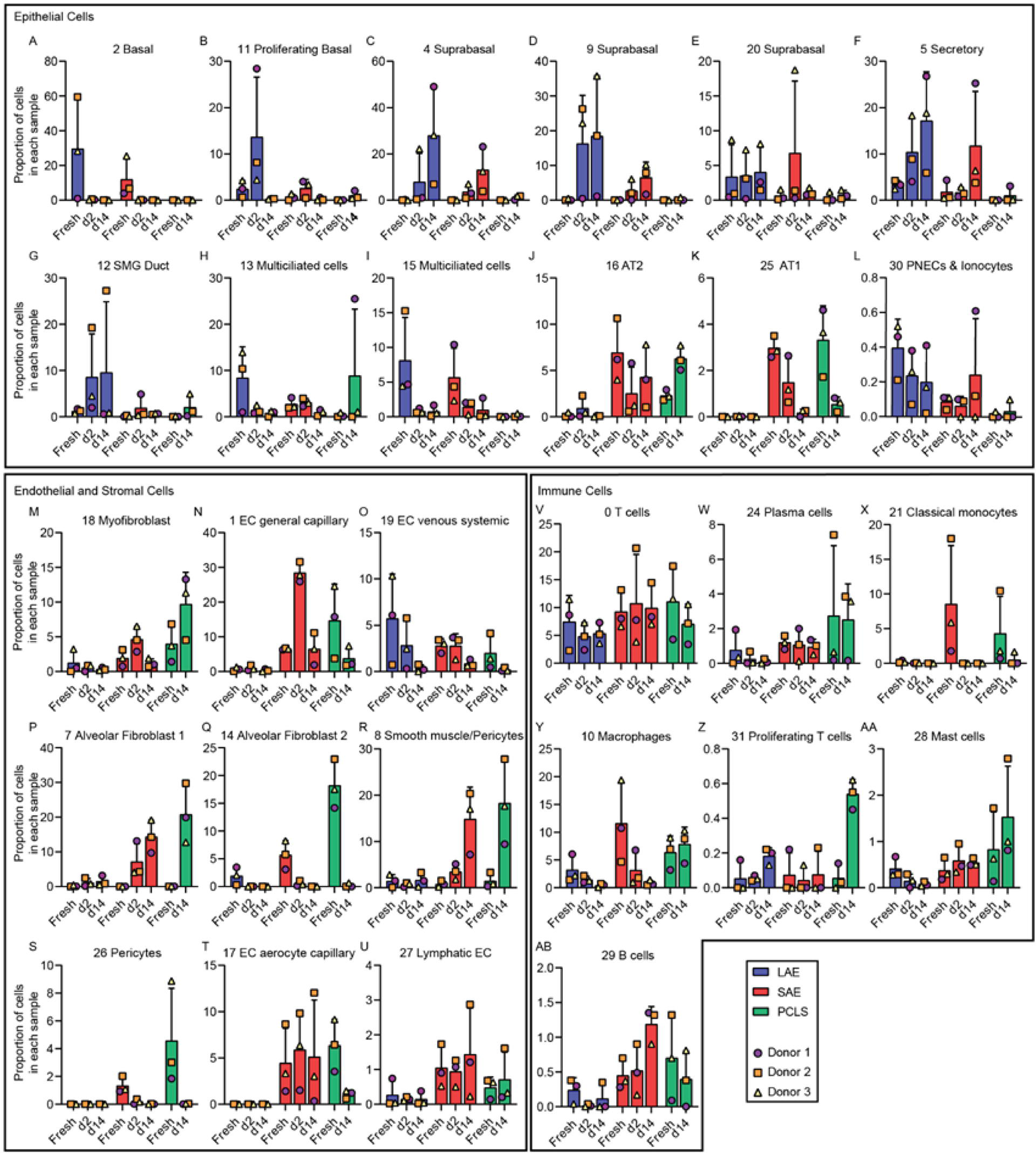
Cell population dynamics over time in the LAE, SAE, and PCLS model. Proportion of A) cluster 2, basal cells B) cluster 11, proliferating basal cells, C) cluster 4, suprabasal cells D) cluster 9, suprabasal cells, E) cluster 20, suprabasal cells, F) cluster 5, secretory cells, G) cluster 12, SMG duct cells, H) cluster 13, multiciliated cells, I) cluster 15, multiciliated cells, J) cluster 16, AT2 cells, K) cluster 25, AT1 cells, L) cluster 30, PNECs and ionocytes, M) cluster 18, myofibroblasts, N) cluster 1, EC general capillary cells, O) cluster 19, EC venous systemic cells, P) cluster 7, alveolar fibroblast 1 (AF1) Q) cluster 14, alveolar fibroblast 2 (AF2) R) cluster 8, smooth muscle/pericytes S) cluster 26, pericytes T) cluster 17, EC aerocyte capillary cells U) cluster 27, lymphatic ECs, V) cluster 0, T cells, W) cluster 24, plasma cells, X) cluster 21, classical monocytes, Y) cluster 10, macrophages, Z) cluster 31, proliferating T cells, AA) cluster 28, mast cells, and AB) cluster 29, B cells in each scRNA-seq sample.

The proportion of secretory cells (cluster 5) in LAE and SAE models as well as SMG duct cells (cluster 12) in the LAE model increased over time (Fig. 4F-G), whereas the proportion of multiciliated cells (cluster 13 and 15) decreased over time in both models (Fig. 4H-I). Our previous H&E and immunofluorescent staining for a-tubulin (Fig. 1C, E, G, and K) demonstrated a well-ciliated epithelium at d14. Thus, we hypothesized that the proportional decrease in multiciliated cells was due to an increase in other cell types rather than ciliated cell loss over time. Indeed, expression of the ciliated cell genes *DNAH11* and *FOXJ1* remained constant in both clusters 13 and 15 (Extended Data Fig. 4J-K) and quantitation of a-tubulin signal confirmed that ciliated cells are retained over 14 days in culture (Extended Data Fig. 4L-Q). AT2 cells (cluster 16) increased in proportion in the PCLS model whereas AT1 cells (cluster 25) decreased over time in both PCLS and SAE models (Fig. 4J-K). PNECs and ionocytes (cluster 30) were low abundance and remained relatively constant over time (Fig. 4L).

With respect to endothelial and stromal cell populations, PCLS exhibited an increase in myofibroblasts (cluster 18; Fig. 4M). General capillary ECs (cluster 1) transiently increased in SAE models at d2 and decreased in PCLS over time (Fig. 4N) whereas venous systemic ECs (cluster 19) decreased over time in all three models (Fig. 4O). The proportion of AF1 (cluster 7) and a subset of pericytes (cluster 8) increased over time whereas AF2 (cluster 14) and a different subset of pericytes (cluster 26) decreased over time in SAE and PCLS (Fig. 4P-S). Aerocyte capillary ECs (cluster 17) were absent in the LAE model and decreased over time in the PCLS (Fig. 4T) whereas lymphatic ECs were relatively stable in all models (Fig. 4U).

With regards to immune cell dynamics, the proportion of T cells (clusters 0) and plasma cells (cluster 24) were constant over time across all models (Fig. 4V-W) while classical monocytes (cluster 21) were only present in fresh tissues (Fig. 4X). The proportion of macrophages (cluster 10) decreased over time in the LAE and SAE models but was maintained in the PCLS model (Fig. 4Y), consistent with replenishment by the proliferating macrophages seen in Fig. 2G. Proliferating T cells (cluster 31) also increased in the PCLS model (Fig. 4Z), again aligning with IF staining (Fig. 2H). B cells and mast cells were present in low numbers and the proportion remained relatively stable over time (Fig. 4AA-AB).

*Explant culture methods extend to animal airway tissue.* To determine whether methods developed for human explants are applicable to animal models, mouse, rabbit and pig tracheal explants were excised from acutely euthanized animals. Histological examination of mouse, rabbit, and pig tracheal explants after 0 days (Extended Data Fig. 5A-C) or 14 days (Fig. 5A-C) in culture revealed well-ciliated epithelia. The electrophysiology of rabbit tracheal explants was investigated after 7, 14, and 21 days in culture (Fig. 5D-J), and dramatically higher responses to Amil, FSK, and CFTRinh-172 in cultured explants were observed compared to previously published measurements of fresh tissue^21^. ENaC activity dropped over time (Fig. 5F), however all other ion transport was preserved (Fig. 5E, G-J). Comparisons of tracheal explants from wildtype and CFTR knockout rabbits^21^ revealed the expected decrease in FSK and CFTRinh-172 response in CFTR knockout tissues (Extended Data Fig. 5D-J). We also characterized the electrophysiology of pig tracheal explants after 7, 14, and 21 days in culture (Extended Data Fig. 5K-Q). In the porcine model, there were trends towards reduced ion transport at d21 compared to d7 measurements with a significant difference in the bumetanide response only.

**Figure 5:**
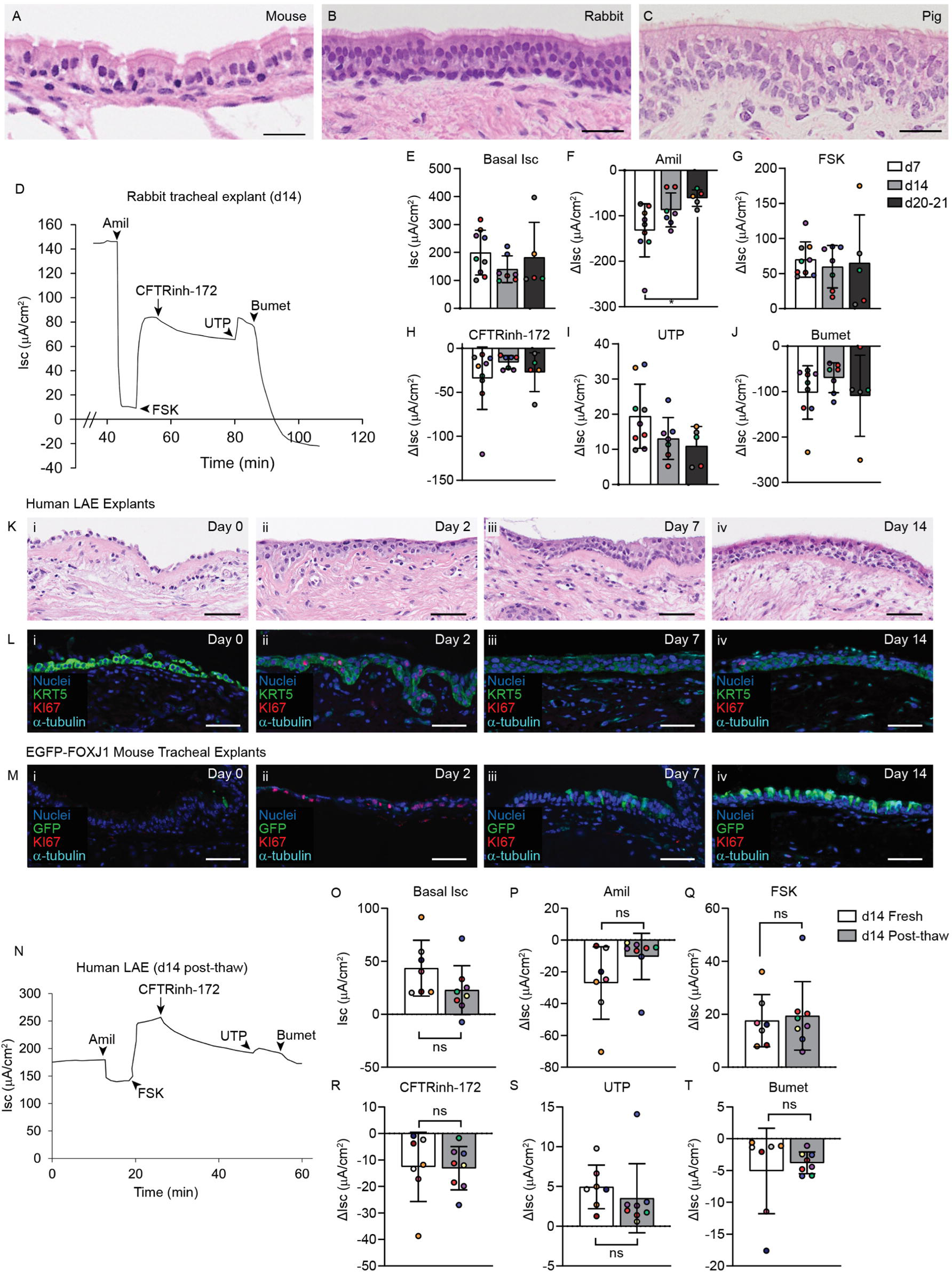
Airway explants can be made from mouse, rabbit, pig, and cryopreserved tissue. A-C) H&E histology of d14 A) mouse, B) rabbit, and C) pig tracheal explants. Scale bars = 25 µm. Representative of N = 3 mice, 3 rabbits, and 4 pigs. D) Representative Ussing tracing of a d14 rabbit tracheal explant. E-J) Time course of rabbit tracheal explant electrophysiology at d7, d14, and d20-21. E) Basal Isc and change in ΔIsc in response to F) Amil, G) FSK, H) CFTRinh-172, I) UTP, and J) Bumet. N = 4-6 animals (represented by different colored dots); 1-2 replicates per animal. One-way analysis of variance (ANOVA) with Tukey’s post-test. K) H&E histology of cryopreserved/thawed human LAE explants at i) 0, ii) 2, iii) 7, and iv) 14 days post-thaw. Scale bars = 50 µm. Representative of N = 2 donors. L) Immunofluorescent localization of KRT5, KI67, and a-tubulin in cryopreserved/thawed human LAE explants at i) d0, ii) d2, iii) d7, and iv) d14 post-thaw. Scale bars = 50 µm. Representative of N = 2 mice. M) Immunofluorescent localization of EGFP-FOXJ1 mouse tracheal explants made from cryopreserved tissue at i) 0, ii) 2, iii) 7, and iv) 14 days post-thaw. Scale bars = 50 µm. N) Representative Ussing tracing of a human LAE explants made from cryopreserved tissue at d14 post-thaw. O-T) Electrophysiology of human LAE explants made from fresh tissue at d14 or cryopreserved tissue at d13-14 post-thaw. O) Basal Isc. Change in ΔIsc in response to P) Amil, Q) FSK, R) CFTRinh-172, S) UTP, and T) Bumet. N = 5-6 donors (represented by different color dots); 1-2 replicates per donor. Unpaired t-test; ns = non-significant.

Finally, we measured mucociliary clearance (MCC) in rabbit tracheal explants by tracking fluorescently labeled beads aerosolized onto the tissue explant surface (Supplemental Video). MCC was higher in explants at all time points than in previously reported *in situ* measurements^22^ (Extended Data Fig. 6A). This finding may indicate that rabbit tracheal explants have more cilia per unit area than the in vivo rabbit trachea, reflecting ∼30% shrinking of explant length upon removal from the animal.

*Airway tissue explants created from cryopreserved tissue.* A challenge of the human airway explant model is dependence on the availability of human tissue. Cryopreservation of human airway tissue would allow investigators to store freshly dissected airway tissues for use on demand. To determine if viable airway explants could be prepared from cryopreserved tissue, freshly excised human bronchial segments were frozen using the CryoStor commercial freezing solution and stored in liquid nitrogen for days-months. Upon thawing, tissues were rinsed in PBS, dissected as previously, and placed on Gelfoam® for culture.

Frozen/thawed LAE exhibited a sparse basal cell layer immediately after thawing, but epithelial cells covered the surface by d2 post-thaw (Fig. 5K). At d7, the thawed epithelium was pseudostratified but poorly ciliated, with cilia returning by d14. IF staining confirmed that the epithelial layer was nearly 100% KRT5+ basal cells at day 0, which proliferated at d2 and differentiated into a-tubulin+ ciliated cells by d14 (Fig. 5L). To further investigate the kinetics of cell renewal, airway explants were created from cryopreserved tracheal tissues from transgenic mice carrying an enhanced green fluorescent protein (EGFP) tag under the FOXJ1 promoter (EGFP-FOXJ1)^23^. In this model, FOXJ1+ ciliated cells are marked by EGFP expression. EGFP expression was lost in the first two days after thawing, but re-emerged by d7, following increased KI67 expression in basal cells at d2 (Fig. 5M). From these data, we conclude that ciliated cells are lost upon airway cryopreservation, but regenerate from basal cells during culture on Gelfoam®. Finally, the function of frozen/thawed human airway explants was assessed by Ussing chamber measurements at day 14 post-thaw. The electrophysiology of frozen/thawed LAE was similar to never-frozen explants (Fig. 5N-T). This finding was confirmed in frozen/thawed rabbit tracheal explants (Extended Data Fig. 6B–J). Thus, we concluded that airway tissue explants generated from cryopreserved human (Fig. 5K-L, N-T) or animal (Fig. 5M; Extended Data Fig. 6B–J) airway tissues regain key morphologic and electrophysiologic features after 14d in culture.

*Human airway explants model viral tropism and host gene responses.* Study of viral infection in human airway tissue explants permits study in an in vivo-like environment with conserved tissue architecture and cells from diverse lineages. Because ex vivo human tissues have been historically limited in longevity, prior studies required viral inoculation on the same day as tissue harvest^24–26^. The practicality of airway explant models would be greatly expanded if tissues could be infected after days or weeks in culture.

A panel of viruses was studied including a GFP-tagged Sendai virus (SeV-GFP; also called murine parainfluenza virus type 1), a GFP-tagged respiratory syncytial virus (RSV-GFP), and the D614G variant of the SARS-CoV-2 in human LAE after 18-45 days in culture. RSV and SARS-CoV-2 are known to cause respiratory disease in humans^27,28^. Though SeV is host restricted to rodents and does not typically cause human disease, it has been proposed as a potential gene therapy vector and infection has been studied in human bronchial epithelial cells in vitro^29^, but not in human explant models. Robust infection was observed at 4 days post-infection (dpi) with all studied viruses as indicated by IF localization of GFP (Fig. 6A-B) or SARS-CoV-2 nucleocapsid (Fig. 6C). Viral infection was also observed in airway explants generated from cryopreserved tissue (Extended Data Fig. 6K).

**Figure 6:**
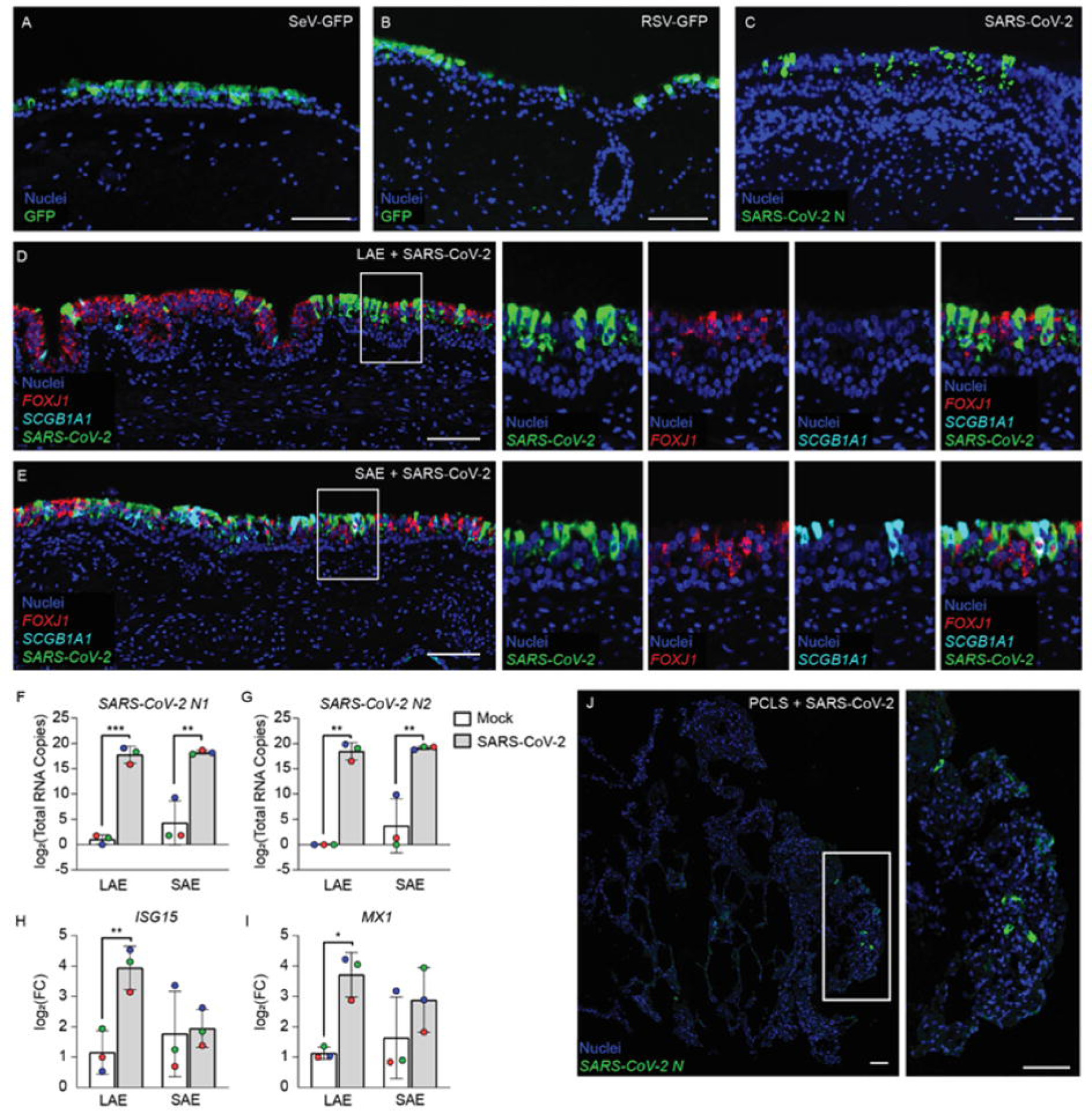
Viral infection in human LAE, SAE, and PCLS models. A-C) Viral infection of human LAE with A) a GFP-tagged Sendai virus (SeV-GFP) after 45 days in culture, B) a GFP-tagged RSV (RSV-GFP) after 45 days in culture, or C) the D614G variant of SARS-CoV-2 after 29 days in culture. Immunofluorescent localization of GFP (A-B) or SARS-CoV-2 (C) at 4 days post-infection (dpi). Scale bars = 100 µm. A-B) Representative of N = 2 donors, 2 replicates/donor. C) Representative of N = 2 donors; 1-2 explants per donor. D-E) Fluorescent RNA *in situ* hybridization of human LAE (D) and SAE (E) inoculated with the D614G variant of SARS-CoV-2 after 25 days in culture. Stained at 4 dpi for *FOXJ1* (red), *SCGB1A1* (cyan), and *SARS-CoV-2* (green). Nuclei counterstained with DAPI. Scale bars = 100 µm. Representative of N = 3 donors. F-G) Log base 2 of the relative copy number of SARS-CoV-2 nucleocapsid genes *N1* (F) and *N2* (G) measured by qRT-PCR and compared to a standard curve in human LAE and SAE explants at 3 dpi. N = 3 donors. H-I) Log base 2 of the relative gene expression of interferon stimulated genes *ISG15* (H) and *MX1* (I) measured by qRT-PCR in human LAE and SAE explants at 3 dpi. The fold change (FC) in gene expression was normalized to *TBP* expression. N = 3 donors. F-I) Data were log transformed before statistical testing due to unequal variances amongst samples and analyzed using a linear mixed-effects model with the donor as a random effect factor; * = p<0.05, ** = p<0.01, *** = p<0.001. J) Representative fluorescent RNA *in situ* hybridization of human PCLS inoculated with the D614G variant of SARS-CoV-2 after 7 days in culture. Scale bars = 100 µm. Representative of N = 4 donors.

More detailed studies focused on the SARS-CoV-2 infection given the importance of the recent COVID-19 pandemic. Human LAE and SAE were inoculated with SARS-CoV-2 and assessed for viral tropism by fluorescent RNA *in situ* hybridization at 4 dpi. As expected, SARS-CoV-2 nucleocapsid transcript was predominantly expressed in FOXJ1+ ciliated cells in both LAE and SAE tissues (Fig. 6D-E). We then quantified RNA copy numbers of SARS-CoV-2 nucleocapsid genes *N1* and *N2* in infected LAE and SAE by performing quantitative real time PCR (Fig. 6F-G). High levels of *N1* and *N2* RNAs were present in both LAE and SAE at 3 dpi but were undetectable in mock-infected LAE and SAE cultures. The interferon-stimulated genes (ISG), *ISG15* and *MX Dynamin Like GTPase 1* (*MX1*), were also elevated at 3 dpi, indicating that airway tissue explants are not only infectable, but can also respond to viral infection by activating interferon-mediated host innate defense pathways (Fig. 6H-I). Finally, to establish an ex vivo infection model in the distal lung, human PCLS tissues were infected with the D614G variant of SARS-CoV-2 after 7 days of culture on Gelfoam®. Infection was less robust in the alveolar epithelium modeled by PCLS compared to LAE and SAE, aligning with previous reports using cultured airway cells and distal lung organoids^30^ (Fig. 6J). From these data, we concluded that human airway explants and PCLS can be used to study viral infection after several weeks in culture.

## Discussion

The Gelfoam®–ALI airway explant model preserves characteristic tissue architecture and diverse cell lineages for at least 14 days in culture. Cell populations within the explant models are dynamic, with certain quiescent cell populations lost (Fig 4A, Q, and S) and repairing cell populations gained (Fig. 4B-D, P, R, and Z). Nonetheless, airway explants provide a unique platform for studying epithelial-stromal and epithelial-immune cell interactions that cannot be studied by standard in vitro methods.

Previous studies have measured airway epithelial electrophysiology immediately after surgical resection^8,17^. Notably, airway explants in our study displayed no measurable electrophysiology when studied immediately after the long cold ischemic time of tissues in this study. However, functional ion transport recovered by d1-2 and d14-15 (Fig. 2I-O) to levels of previously reported fresh tissue measurements^8,17^ indicating a remodeling and/or re-oxygenation of airway explants during the Gelfoam® culture. Indeed, a proliferating basal cell population was detected by scRNA-seq (Fig. 4B) and confirmed by IF in d2 explants (Extended Data Fig. 4E). This population was lost by d14 (Fig. 4B; Extended Data Fig. 4F), consistent with transient repair followed by a return to quiescence. These data highlight the utility of the explant model for studying injury and repair ex vivo.

The recovery of airway tissue function after ∼40h of cold ischemic injury is important, as it greatly increases the number of lungs that can be studied. With time in culture, our scRNA-seq data revealed a subset of basal/suprabasal cells expressing transcripts associated with airway epithelial hypoxia in LAE and SAE explants^20^. Metabolically active basal cells in LAE may partially receive O_2_ from submucosal systemic capillaries, and absence of explant perfusion may contribute to hypoxia. Additionally, airway epithelia gain O_2_ from the lumen, and accumulated mucus may result in O_2_ limitation. While mucin secretion is a key property of airway epithelia for maintaining mucociliary clearance, this observation highlights the importance of removing apical mucus from airway explants to maintain epithelial homeostasis over time. Alternatively, airway explants maintained on Gelfoam® sponge without mucus removal induce asymmetric apical hypoxia to model muco-obstructive lung diseases, including cystic fibrosis, chronic obstructive lung disease, and asthma.

The ability to prepare viable airway explants from cryopreserved tissue presents another major advancement in the field. While cryopreservation of PCLS has been previously reported^31–33^, our study represents the first application of this methodology to generate viable airway tissue explants from cryopreserved tissue. Airway explants from cryopreserved tissue successfully replicated morphologic and electrophysiologic features of fresh tissue. Though cryopreservation severely injures the airway epithelium (Fig. 5L-M), KRT5+ basal cells were retained and regenerated a viable, ciliated epithelium. Future scRNA-seq studies to characterize other cellular changes in the cryopreservation recovery process should be informative for understanding mechanisms of airway regeneration. *In vitro*, airway basal cells can be efficiently transduced by gene transfer vectors for genetic modification^34,35^. Thus, unobstructed access to the basal cell layer following cryo-injury could facilitate gene modulation in ex vivo human tissues, a unique application of this technology.

The ability to culture viable explants from diverse animal models could improve translation between animal and human studies. Further, the ability to generate multiple explants per animal and the ability to cryopreserve excess tissue for later experiments aligns well with the “three Rs principle of animal research”, Replacement, Reduction and Refinement^36^. Unlike in vivo experiments where only one condition can be tested per animal, explant culture allows airway tissue to be divided into multiple explants, e.g., 20 or more explants per adult rabbit trachea, permitting the study of multiple conditions and/or replicates per animal. Another advantage is the ability to create explant cultures from animal tissues carrying genetic modification. This feature was demonstrated here through the use of CFTR knockout rabbits^21^ and EGFP-FOXJ1 transgenic mice^23^.

The Gelfoam®-cultured airway explants can be used to study viral infection with SeV, RSV, and SARS-CoV-2. SARS-CoV-2 tropism in the airway explant was comparable to COVID-19 autopsy samples indicating ciliated cells as the primary target of viral entry in the airways^30^ with lower infection rates in the distal lung^29^. Previous groups have studied viral infection in ex vivo tissue models^24,37^ including PCLS^38,39^. However, all prior work was restricted to short-term time points and required inoculation immediately after tissue procurement. In our study, airway tissue explants cultured on Gelfoam® could be infected after 18-45 days in culture. The ability to infect airway tissue explants after extended culture or following cryopreservation (Extended Data Fig. 6K) greatly expands this model’s practicality. This feature also gives airway tissue explants time to recover from cold ischemia and dissection before viral inoculation. Here, we studied viral infection at 4 dpi, but studies of late-stage recovery from viral infection are possible and are a future application of this work.

The current form of the Gelfoam®–explant has limitations: Rabbit tracheal explant electrophysiology differed from published measurements of fresh rabbit trachea^21^, with significantly higher rates to amiloride, forskolin, and CFTRinh-172 sensitive currents than fresh tissues (Fig. 5D-J). Pig tracheal explants electrophysiology also differed from previously published measurements of freshly excised tissue^40,41^, with a greater basal Isc (∼threefold increase) and a greater forskolin response (∼fivefold increase) in our d14 pig tracheal explants compared to fresh tissue measurements. Future work will be required to identify the mechanisms underlying enhanced ion transport in the rabbit and pig explant model and why these differences were not observed in the human explant model. We also observed that relatively thinner airway explants, e.g., rabbit tracheal explants, exhibited gradual shrinkage from the edge over time in culture (Extended Data Fig. 6L). This observation underscores the need for further optimization of explant culture methods for thinner airway explant tissues.

Further, although the in vivo alveolar space is typically quiescent, high proliferation rates in various cellular compartments were observed in the d14 PCLS model (Fig. 2D-H). This proliferation may be a response to the mechanical injury incurred during PCLS preparation (tissue slicing on a Compresstome®). Alternatively, the prolonged proliferation could point to innate differences in culture requirements or differences in how the airway and alveolar epithelium respond to the material properties and mechanical cues of Gelfoam®. Though Gelfoam®-culture prolonged PCLS viability considerably (Fig. 2C), further media optimization and exploration of new matrix substrata may be warranted.

In summary, we have shown that culture of human airway explants and PCLS at air-liquid interface on a Gelfoam® sponge prolongs tissue viability for several weeks with preservation of diverse organotypic cell types. These methods are generally applicable to animal models including mouse, rabbit, and pig. Airway tissue explants created from cryopreserved tissue mimic fresh explants and airway tissue explants can be used to model viral infection and host gene response. Overall, the Gelfoam®–ALI explant model expands the practicality of ex vivo airway tissue studies and could enable many important future applications.

## Methods

*Preparation of Gelfoam®.* Gelfoam® (Ethicon 1972) was cut using sterile scissors into ∼1cm x 1cm squares and was placed in a six well-plate (Corning #3516). Explant media, prepared by adding L-Glutamine (Gibco #25030-081), penicillin-streptomycin (Gibco #15140-122), and 10% heat inactivated fetal bovine serum (Gibco #16140-071) to Dulbecco’s Modified Eagle Medium, high glucose (DMEM-H; Gibco #11965-092), was dispensed into the plate at 3 mL per well. The Gelfoam® was rehydrated by gently pressing the Gelfoam® sponge into explant media using ethanol-sterilized forceps. A cocktail of supplemental antibiotics and antifungals including amphotericin B (1 μg/mL), ceftazidime (100 μg/mL), tobramycin (80 μg/mL), vancomycin (100 μg/mL), mycamine (20 μg/mL), and diflucan (25 μg/mL) were added during the first 24 hours of culture. Additional antibiotics and/or antifungals were added for CF tissues as before^42^ according to the patient’s clinical records. The explant media surrounding the Gelfoam® was removed and replaced every 2-3 days.

*Airway Tissue Dissection*. Large airway explants (LAE) were obtained from first to third generation bronchi from previously healthy deceased lung donors or from cystic fibrosis (CF) transplant lungs obtained under University of North Carolina’s Office of Human Research Ethics/Institutional Review Board study #03-1396. To prepare LAE, a 1 cm segment of unbranched bronchus was removed and cut longitudinally to expose the airway lumen. The surface epithelium was then stripped away from the underlying cartilage using surgical micro-scissors under a dissecting microscope. Small airway explants (SAE) were obtained from bronchioles with diameter less than 2mm. Bronchioles were microdissected from the surrounding parenchyma as previously described^43^ and cut longitudinally to expose the airway lumen. After dissection, LAE and SAE were placed on a media-soaked Gelfoam® sponge with the lumen exposed to air and maintained in a high humidity incubator. Mouse, rabbit, and pig tracheal explants were prepared by the same methods except that the mouse tracheal epithelium was left attached to the underlying cartilage. Rabbits were anesthetized with isoflurane and exsanguinated by severing the abdominal aorta when they were on an anesthetic plan. Mice were euthanized by CO2 inhalation and pigs were euthanized by barbituric IV injection following sedation. We studied female wildtype and EGFP-FOXJ1 transgenic mice on a mixed C3H × C57BL/6 background^23^, wildtype and CFTR knockout New Zealand White (NZW) rabbits^21^, and wildtype American Yorkshire × Duroc pigs. All animal experiments were approved by the University of North Carolina Institutional Animal Care and Use Committee and performed in accordance with the guidelines and regulations governing the use of these laboratory animals.

*Preparation of precision cut lung slices (PCLS).* Human PCLS were prepared from previously healthy donor lungs as previously described^10,44–46^. Briefly, a piece of distal lung was isolated, warmed to room temperature, and inflated using a needle and syringe with warm (42°C) 3% low-melting point agarose (ThermoFisher #BP165-25) diluted in DMEM-H (Gibco #11965-092). The inflated piece of lung was then cooled to 4°C for at least 30 minutes to solidify the agarose. An Acu-punch Biopsy Punch (ThermoFisher #NC9324386) was used to cut 8 mm cores from the inflated lung. Cores were glued to the end of a specimen tube plunger using Scotch super glue (3M #AD119) and loaded onto a Compresstome® (Precisionary) where the tissue was cut to 300 µm thickness. PCLS were then placed on Gelfoam® for long-term culture and monitored using a WST-8 cell counting kit (VWR #89155-898) according to the kit’s instructions. Briefly, PCLS were moved to a 24 well plate using forceps and submerged in 500 uL WST-8 assay media for 1h at 37°C. The PCLS were then returned to Gelfoam® or submerged culture conditions and samples of the WST-8 assay media were transferred to a 96-well assay plate for colorimetric reading using a FLUOstar Omega microplate reader (BMG LABTECH).

*Measurements of Isc by Ussing chamber.* Isc recordings in Ussing chambers were conducted on human LAE and SAE, rabbit tracheal, and pig tracheal explants according to previously published protocols^21,47^. Human, rabbit, and pig airway tissues were mounted with an exposed surface area of 0.025 cm^2^, 0.094 cm^2^, and 0.053 cm^2^, respectively. Both sides of the airway explant were perfused with equal amounts (10 mL) of standard Krebs Bicarbonate Ringer’s (KBR) solution (140 Na^+^ mM, 120 Cl^-^ mM, 5.2 K^+^ mM, 1.2 Mg^2+^ mM, 1.2 Ca^2+^ mM, 2.4 HPO_4_^2-^ mM, 0.4 H_2_PO_4_^-^ mM, and 25 HCO_3_^-^ mM) at 37°C and gassed with 95% O_2_ and 5% CO_2_, providing gas lift circulation. Samples were equilibrated under voltage clamp (Isc) conditions for at least 20 minutes until a baseline Isc was achieved. Drugs were added as follows: amiloride (apical, 10 µM), forskolin (basolateral, 10 µM), CFTRinhibitor-172 or GlyH-101 as indicated (apical, 10 µM), uridine-5’-triphosphate (UTP) (apical, 100 µM), and bumetanide (basolateral, 1 mM).

*Mucociliary clearance*. To measure mucociliary clearance, fluorescently labeled beads (200 nm) were aerosolized onto the surface of airway tissue explants, visualized under a dissecting microscope, and recorded, as previously^22^.

*Immunostaining and RNA in situ hybridization.* Explant tissues were fixed by submersion in 10% neutral buffered formalin (NBF) for 2-7 days, washed using PBS and stored in 70% ethanol before embedding and serial sectioning. Formalin-fixed paraffin-embedded (FFPE) sections were stained with hematoxylin and eosin (H&E) and Alcian Blue Periodic acid Schiff (AB-PAS). FFPE sections were also assessed by immunofluorescent localization as previously described^48^ (see Extended Data Table 1 for antibodies and dilutions). Colorimetric or fluorescent RNA *in situ* hybridization was performed as previously described^43^ (see Extended Data Table 2 for a list of materials).

*Cryopreservation and thawing of airway tissue explants.* Small (∼1 cm) segments of bronchus or microdissected bronchioles were isolated and transferred to cryovials containing 1 mL CryoStor freezing medium (Millipore Sigma #C2999) as intact airways. Cryovials were stored in a Corning CoolCell Freezing Container (Corning #432000) and subsequently frozen at -80°C. After 48-72 hours, cryovials were then transferred to liquid nitrogen for longer term storage. To thaw, cryovials were incubated in a 37°C water bath until all CryoStor freezing medium was melted. Airway tissue was then rinsed in PBS, cut longitudinally to expose the airway lumen, and placed on a Gelfoam® sponge for long-term culture. PCLS tissue cut using a Compresstome® were frozen and thawed in the same manner.

*Single-cell dissociation of LAE, SAE, and PCLS tissues.* LAE, SAE, and PCLS tissues were cryopreserved in CryoStor freezing medium after 0 (fresh), 2, and 14 days in culture. Cryovials were thawed, rinsed in PBS, placed into a 15 mL tube containing 5 mL of Accutase solution (Sigma-Aldrich, #A6964-100ML) with EDTA (invitrogen, #15575-038) (5 mM) and pronase (Roche, #10165921001) (1 mg/mL), and incubated on a shaker for 30 min at 37°C. Following incubation, the tube was centrifuged (600 g, 2 min, 4 °C), the supernatant was discarded, and 5 mL of Hank’s Balanced Salt Solution (Ca-, Mg-) (gibco) buffer containing liberase (Roche, #05401119001) (50 mg/mL) and DNase (Roche, #10104159001) (25 ug/mL) was added to the tube. The sample was then incubated on the shaker for 10 min at 37°C. After incubation, the cell suspension was gently pipetted 20 times using a wide-orifice pipette tip. To neutralize enzymes, 500 mL of fetal bovine serum was added to the cell suspension, and the neutralized mixture was filtered through a 100 um filter, followed by centrifugation (600 g, 2 min, 4 °C). After removing supernatant, dissociated cells were resuspended in PBS (Ca-, Mg-) containing 0.01% Ultrapure BSA (Invitrogen, #AM2616), filtered through a 40-μm cell strainer (Bel-Art, H13680-0040), and centrifuged (600 g, 2 min, 4 °C). Cell count and viability was assessed using trypan blue dye exclusion with a Countess 3 automated cell counter (Thermo Fisher Scientific), and the cell concentration was adjusted to approximately 1000 cells/uL. The dissociated cells were used for 10X Genomics scRNA-seq library preparation.

*Chromium 10X Genomics scRNA-seq library preparation and sequencing*. Single cells were captured using a 10X Chromium controller (10X Genomics) and libraries were prepared following the Single Cell 3’ Reagent Kits v3.1 (Dual Index) User Guide (10X Genomics). Cellular suspensions were loaded on a Chromium Controller instrument (10X Genomics) to generate single-cell Gel Bead-In-EMulsions (GEMs). Reverse transcription (RT) was performed in a Veriti 96-well thermal cycler (ThermoFisher). After RT, GEMs were harvested, and the cDNA underwent size selection with SPRIselect Reagent Kit (Beckman Coulter). Indexed sequencing libraries were constructed using the Chromium Single-Cell 3’ Library Kit (10X Genomics) for enzymatic fragmentation, end-repair, A-tailing, adapter ligation, ligation cleanup, sample index PCR, and PCR cleanup. Barcoded sequencing libraries were quantified by Agilent Tapestation. The libraries were loaded onto a NovaSeq 6000 (Illumina) with a custom sequencing setting of paired-end reads (28 and 90 cycles for Read 1 and 2, respectively), to achieve a sequencing depth of at least ∼5 x 10^4^ reads per cell.

*scRNA-seq data analysis*. Sequencing data in FASTQ format per sample were processed and mapped to recent reference human genome and Gencode gene annotation using Cell Ranger v7 pipeline from 10xGenomics. The resultant gene x cell matrices were imported into R using the *Seurat* package^49^ after ambient RNA contamination adjustments using R package, *SoupX*^50^ with recommended parameters. Cells were filtered at >300 gene counts, and maximal gene counts varying between 7000-9000 gene counts, and <20% mitochondrial gene expression. The samples were integrated into a combined data set using the IntegrateData function from *Seurat* with targeted 4000 anchor genes and 50 PCA dimensions. Cell clusters from integrated data set were generated with the FindClusters function with 40 PCA dimensions and resolution parameter of 0.5. Cell type predictions were made using local setup of *azimuth* (https://azimuth.hubmapconsortium.org/) against the core human lung cell atlas (HLCA) version 2 from normal lung samples^18^. Final cell type labels were assigned manually based on marker genes analysis with FindAllMarkers function from Seurat and predicted cell types. Various plots were generated with Seurat and the R package, *ggplot2* (Wickham H (2016). *ggplot2: Elegant Graphics for Data Analysis*. Springer-Verlag New York. ISBN 978-3-319-24277-4).

*Viral inoculation.* Airway tissue explants and PCLS were removed from the Gelfoam® sponge and transferred to an empty well of a 12-well plate (Corning #3513) using sterile forceps. Explants were then submerged in a virus-containing inoculum and incubated for 2 hours at 37°C. For SARS-CoV-2 inoculation, samples were submerged in 300 uL virus (1.91x10^7^ PFU) in a 24 well plate. Samples were submerged in 100 uL virus in a 48 well plate for Sendai virus (0.8Log10 TCID50/ml) and RSV (0.65Log10 TCID50/ml) inoculations. After a PBS rinse, explants were transferred back to the Gelfoam® sponge and cultured for 3-4 days. Gene expression of *interferon stimulated gene 15 (ISG15), MX Dynamin Like GTPase 1 (MX1), and* SARS-CoV-2 nucleocapsid genes, *N1* and *N2*, was assessed by quantitative real time polymerase chain reaction (qRT-PCR). For *N1* and *N2*, the Ct values were compared to a standard curve to obtain the viral copy number.

*Statistics.* All data were analyzed using GraphPad Prism 9.5 (RRID:SCR_002798) by an unpaired t-test for single comparisons or a one-way analysis of variance (ANOVA) with Tukey’s or Dunnett’s post-test for multiple comparisons, as indicated in the figure legends. All data are presented as mean ± standard deviation (SD).

## Acknowledgements

The authors thank the UNC Marsico Lung Institute Tissue Procurement and Cell Culture Core for providing human lung tissue and the UNC Histology Research Core. We gratefully acknowledge Dr. Larry Ostrowski for provision of wildtype and FOXJ1-EGFP transgenic mice. This project was supported in part by the National Institutes of Health (1T32GM133364 and 5F31HL158197 to R.E.L.; R01HL163602 to K.O.; 1K12TR004416 to S.A.S.; 1R21AI138247 to R.J.P.; and 2P30DK065988 to R.C.B.), the Cystic Fibrosis Foundation (RANDEL17XX0 and RANDEL20XX2 to S.H.R; GRUBB2110 to B.R.G.; PICKLE18I0 to R.J.P; BOUCHE19XX0 to R.C.B.; BOUCHE22G0-COLLAB to R.C.B. and K.O.; OKUDA20G0 to K.O.), and a research grant from the Cystic Fibrosis Research Institute to K.O. This project was also partially supported by the Rapidly Emerging Antiviral Drug Development Initiative (READI) at the University of North Carolina at Chapel Hill, funded by the North Carolina coronavirus state and local fiscal recovery funds program appropriated by the North Carolina General Assembly.

## Author contributions

R.E.L., K.O., S.H.R., and R.C.B. conceived the project and, with R.J.P, J.S.H., R.S.B., W.K.O., and B.R.G., designed the experiments. R.E.L., K.O., J.R.G., S.A.S., and S.C.G. dissected and cultured LAE and SAE explants. J.R.G. and L.R. prepared PCLS. R.E.L. and S.A.S. performed immunostaining and R.G. and K.O. performed RNA in situ hybridization. J.R.G. performed WST-8 viability assays. B.R.G and T.D.R performed Ussing chamber measurements and mucociliary clearance measurements. R.E.L dissected mouse tracheal explants; B.R.G and T.D.R dissected rabbit tracheal explant; and E.C. resected and provided porcine lung tissue. Single cell RNA-sequencing sample preparation was performed by R.E.L., J.R.G., and K.O., and H.D., K.O., S.A.S., A.M.C., and R.E.L performed the subsequent data analysis. Viral inoculations were performed by R.E.L. and C.E., and S.N. performed qPCR. R.E.L. drafted the manuscript and composed all figures. All authors read, edited, and approved the final version of the manuscript.

