## Extended Data for "Prolonged airway explant culture enables study of health, disease, and viral pathogenesis"

*Supplemental Table 1:* *Formalin fixed paraffin embedded (FFPE) immunohistochemistry protocol.* Primary and secondary antibodies along with antibody source, catalog number, concentration, and RRID.

| **Primary Antibody Target** | **Source** | **Catalog number** | **Concentration** | **RRID** |
| --- | --- | --- | --- | --- |
| a-Tubulin | Millipore | MAB1864 | 3 µg/ml | RRID:AB_2210391 |
| MUC5B | Sigma-Aldrich | HPA008246 | 0.1 µg/ml | RRID:AB_1854203 |
| MUC5AC | Thermo Fisher Scientific | MA5-12178 | 4 µg/ml | RRID:AB_10978001 |
| PECAM1 | AbCam | ab76533 | 1.3 µg/ml | RRID:AB_1523298 |
| ACTA2 | LSBio (LifeSpan) | B3933 | 1 µg/ml | RRID:AB_10686658 |
| PTPRC | Thermo Fisher Scientific | 14-0459-82 | 1 µg/ml | RRID:AB_467274 |
| CD68 | Cell Signaling Technology | 76437 | 1.5 µg/ml | RRID:AB_2799882 |
| CD3 | AbCam | ab16669 | 1:100 | RRID:AB_443425 |
| CD20 | Agilent | M0755 | 1:200 | RRID:AB_2282030 |
| RAGE/AGER | R&D Systems | AF1145 | 0.4 µg/ml | RRID:AB_354628 |
| Pro-SPB | Seven Hills Bioreagents | WRAB-55522 | 1:1000 | RRID:AB_2938816 |
| LAMP-3 | Novus | DDX0191P-100 | 5 µg/ml | RRID:AB_2827532 |
| Ki-67 | BD Biosciences | 550609 | 1 µg/ml | RRID:AB_393778 |
| PDGFRA | R&D Systems | AF-307-NA | 1:300 | RRID:AB_354459 |
| Keratin 5 | Biolegend | 905901 | 2 µg/ml | RRID:AB_2565054 |
| GFP | AbCam | ab6556 | 0.5 µg/ml | RRID:AB_305564 |
| SARS-CoV-2 N | Thermo Fisher Scientific | PA1-41098 | 1 µg/ml | RRID:AB_1087200 |
| **Secondary Antibodies** | **Source** | **Catalog number** | **Concentration** | **RRID** |
| Donkey anti-mouse 488 | Jackson ImmunoResearch Labs | 715-545-151 | 3.125 µg/ml | RRID:AB_2341099 |
| Donkey anti-rabbit 488 | Jackson ImmunoResearch Labs | 711-545-152 | 3.125 µg/ml | RRID:AB_2313584 |
| Donkey anti-goat 488 | Jackson ImmunoResearch Labs | 705-545-147 | 3.125 µg/ml | RRID:AB_2336933 |
| Donkey anti-rabbit 555 | Thermo Fisher Scientific | A31572 | 2 µg/ml | RRID:AB_162543 |
| Donkey anti-rabbit 594 | Jackson ImmunoResearch Labs | 711-585-152 | 3.125 µg/ml | RRID:AB_2340621 |
| Donkey anti-rat 594 | Thermo Fisher Scientific | A21209 | 2 µg/ml | RRID:AB_2535795 |
| Donkey anti-mouse 647 | Thermo Fisher Scientific | A31571 | 2 µg/ml | RRID:AB_162542 |
| Donkey anti-goat 647 | Thermo Fisher Scientific | A21447 | 2 µg/ml | RRID:AB_2535864 |
| Donkey anti-chicken 647 | Jackson ImmunoResearch Labs | 703-605-155 | 3.125 µg/ml | RRID:AB_2340379 |

*Supplemental Table 2:* *Formalin fixed paraffin embedded (FFPE) RNA in situ hybridization protocol.* RNA in situ hybridization probes along with probe source, catalog number, concentration, and RRID.

| **Colorimetric RNA in situ hybridization probe** | **Source** | **Catalog number** |
| --- | --- | --- |
| RNAscope™ Probe- Hs-FOXJ1 | Advanced Cell Diagnostics, Inc. | 430921 |
| RNAscope™ Probe- Hs-SCGB1A1 | Advanced Cell Diagnostics, Inc. | 469971 |
| RNAscope™ Probe- Hs-KRT5-O1 | Advanced Cell Diagnostics, Inc. | 547901 |
| RNAscope™ Probe- Hs-CFTR | Advanced Cell Diagnostics, Inc. | 603291 |
| RNAscope™ Probe- Hs-CD68 | Advanced Cell Diagnostics, Inc. | 560591 |
| RNAscope™ Probe- Hs-SFTPB | Advanced Cell Diagnostics, Inc. | 544251 |
| RNAscope™ Probe- Hs-AGER | Advanced Cell Diagnostics, Inc. | 470121 |
| **Fluorescent RNA in situ hybridization probes** |  |  |
| RNAscope™ Probe- Hs-SCGB1A1 | Advanced Cell Diagnostics, Inc. | 469971 |
| RNAscope™ Probe- Hs-FOXJ1-C2 | Advanced Cell Diagnostics, Inc. | 430921-C2 |
| RNAscope™ Probe- V-nCoV2019-S-C3 | Advanced Cell Diagnostics, Inc. | 848561-C3 |
| RNAscope™ Probe- Hs-SFTPC | Advanced Cell Diagnostics, Inc. | 452561 |
| RNAscope™ Probe- Hs-CFTR-C2 | Advanced Cell Diagnostics, Inc. | 603291-C2 |
| RNAscope™ Probe- Hs-SCGB1A1-C3 | Advanced Cell Diagnostics, Inc. | 469971-C3 |

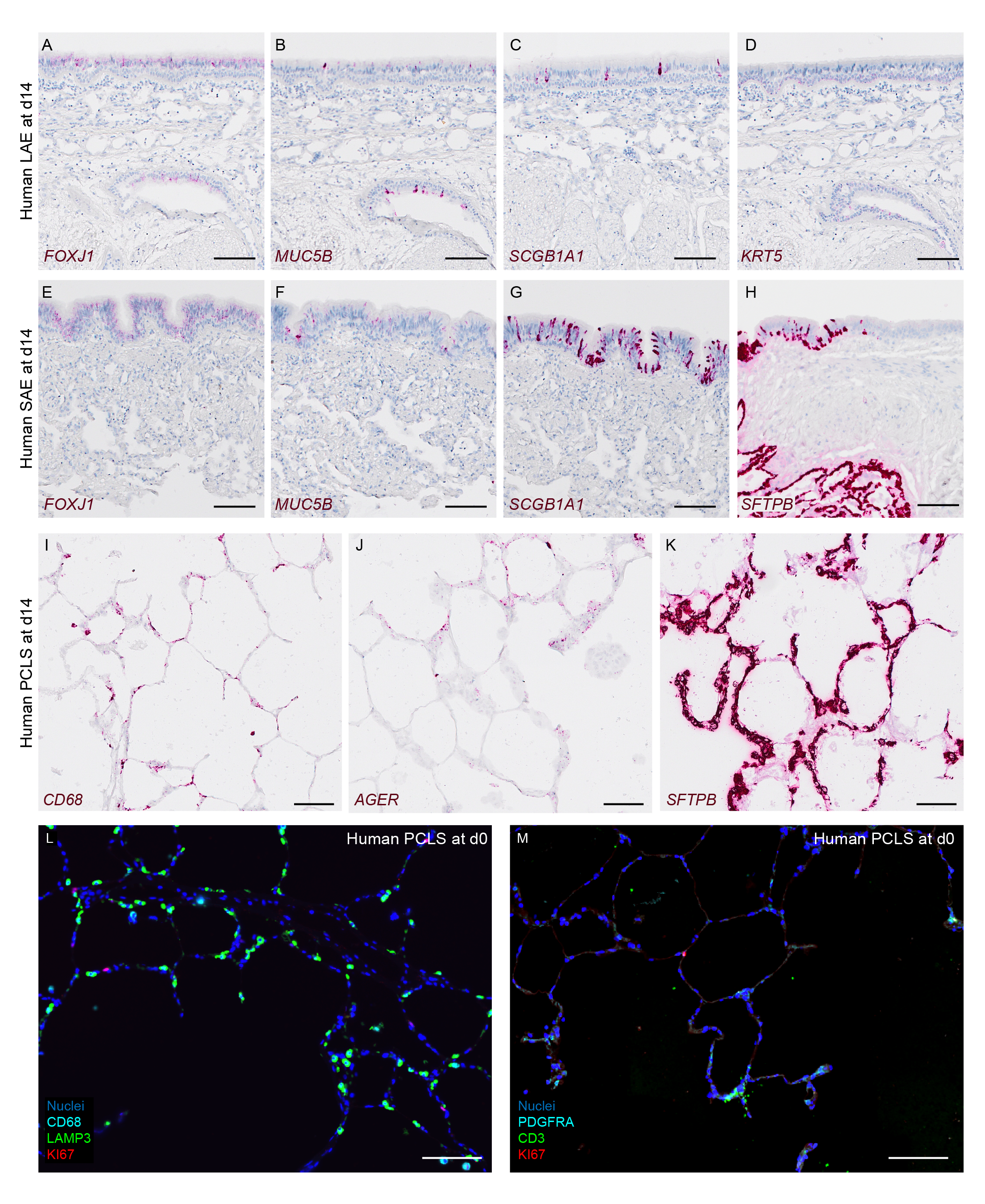

*Extended Data Fig 1. Human airway explants and PCLS express characteristic markers of epithelial, endothelial, and immune cell populations.* A-H) Representative colorimetric RNA *in situ* hybridization images. *FOXJ1* transcript in d14 LAE (A) and SAE (B) explants. *MUC5B* transcript in d14 LAE (C) and SAE (D) explants. *SCGB1A1* transcript in d14 LAE (E) and SAE (F) explants. G) *KRT5* transcript in a d14 LAE explant. H) *SFTPB* transcript in a d14 SAE explant. I-K) Representative colorimetric RNA *in situ* hybridization for I) *CD68*, J) *AGER*, and K) *SFTPB*. L-M) Immunostaining of d0 PCLS for CD68, LAMP3, and Ki67 (L) and PDGFRA, CD3, and Ki67 (M). Representative of N = 3 donors. All scale bars = 100 µm.

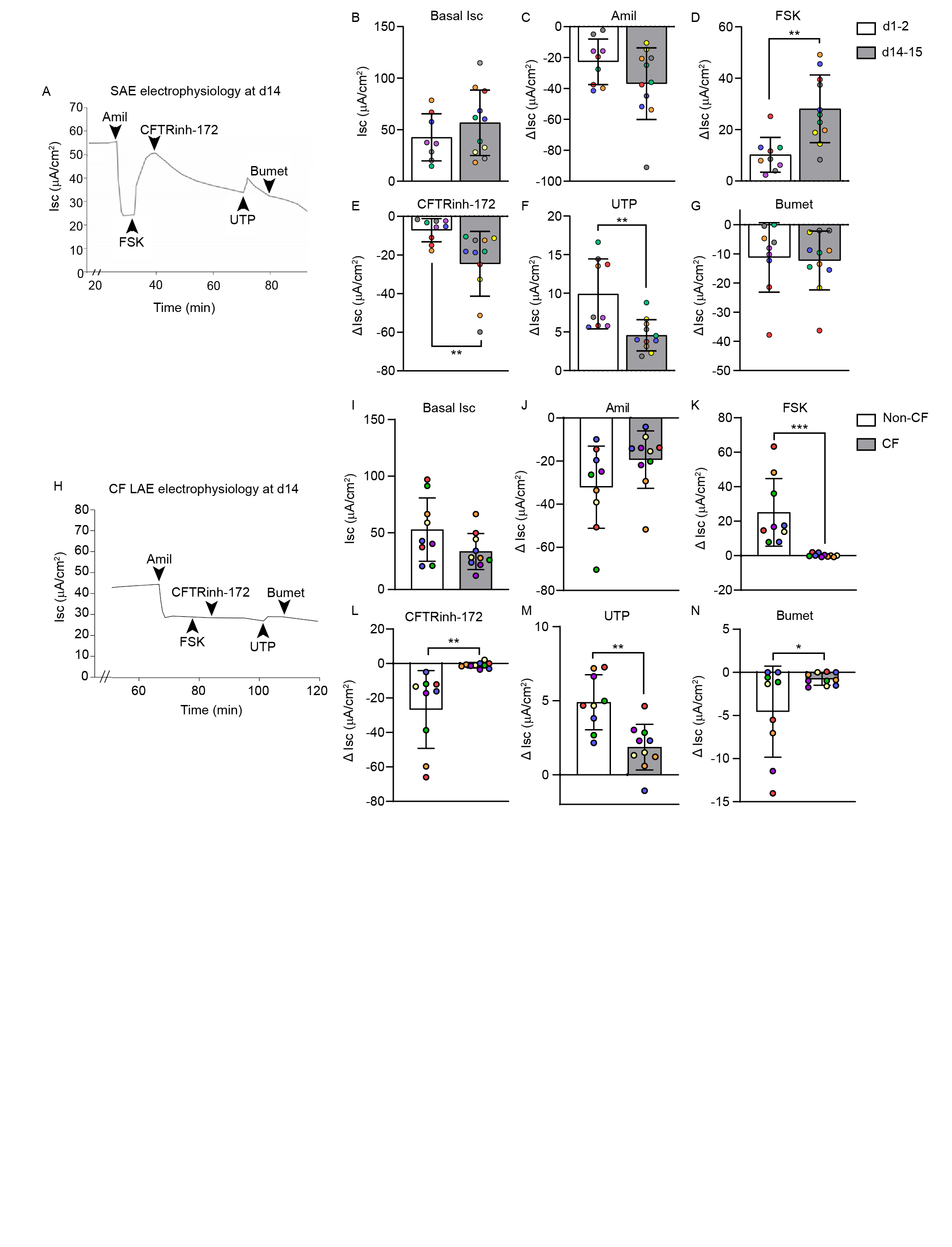

*Extended Data Fig 2. Electrophysiology of SAE explants and CF versus non-CF LAE explants.* A-G) Electrophysiology of human LAE explants at d1-2 and d14-15. A) Representative Ussing tracing of a d14 human SAE explant. B) Basal short circuit current (Isc) and ΔIsc in response to C) amiloride (Amil), D) FSK, E) CFTRinh-172, F) UTP, and G) bumetanide (Bumet). H-N) Electrophysiology of human CF vs non-CF LAE explants at d14. H) Representative Ussing tracing of a CF LAE explant at d14. I) Basal Isc and ΔIsc in response to J) Amil, K) FSK, L) CFTRinh-172, M) UTP, and N) Bumet.

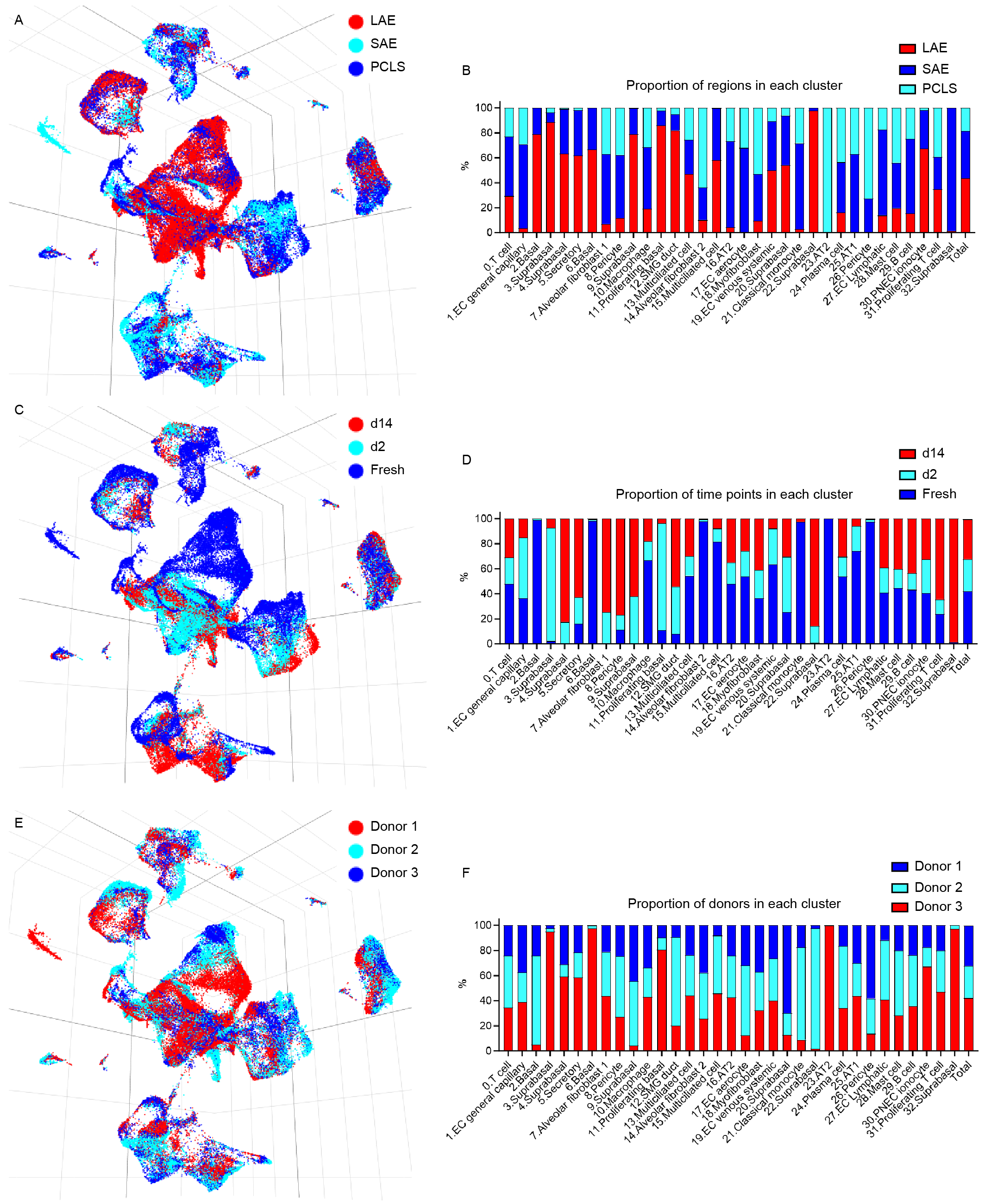

*Extended Data Fig 3. Single cell RNA-sequencing data shown by region (LAE, SAE, PCLS), time point, and cell donor.* A) UMAP colored by region. B) Proportion of the regions that makes up each cluster. C) UMAP colored by time point. D) Proportion of the time points that makes up each cluster. E) UMAP colored by cell donor. F) Proportion of the cell donors that makes up each cluster.

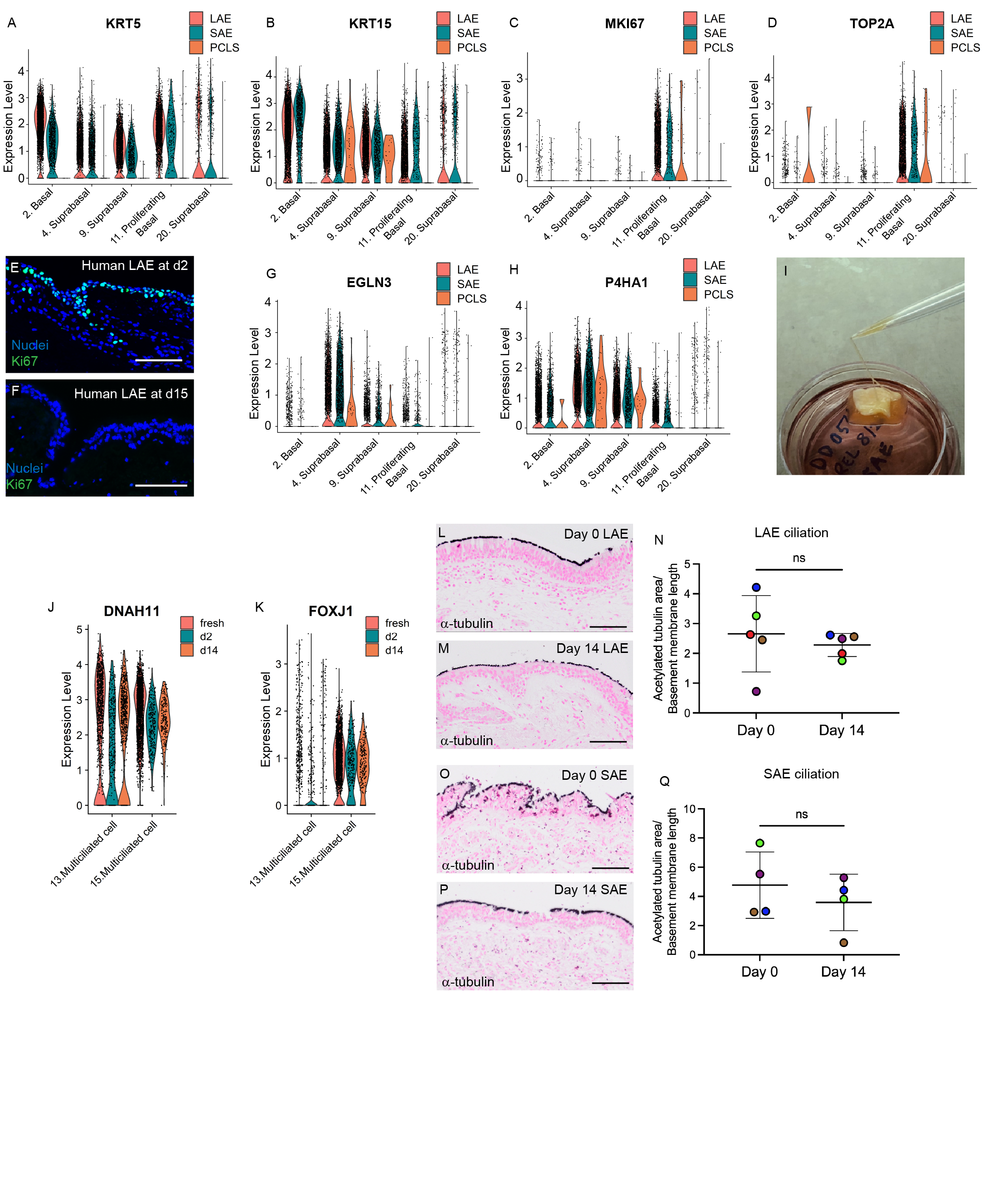

*Extended Data Fig 4. Basal and suprabasal clusters in the LAE and SAE models.* A-D) Violin plot of the basal cell markers, *KRT5* (A) and *KRT15* (B) and the proliferation markers, *MKI67* (C), and *TOP2A* (D) in all basal and suprabasal clusters. E-F) Immunofluorescence for KI67 in human LAE at d2 (E), and d15 (F). G-H) Violin plot of the hypoxia markers, *EGLN3* (G) and *P4HA1* (H) in all basal and suprabasal clusters. I) Mucus harvested from the apical surface of a human LAE at d28. L-J) Violin plot of the ciliated cell markers, *DNAH11* (J) and *FOXJ1* (K) in all multiciliated cell clusters. L-M) Immunohistochemistry for a-tubulin to mark ciliated cells at d0 (L) and d14 (M). Representative of N = 5 donors. N) LAE quantitation of a-tubulin area normalized to basement membrane length. N = 5 donors (represented by different colored dots). Paired t-test; ns = non-significant. O-P) Immunohistochemistry for a-tubulin to mark ciliated cells at d0 (O) and d14 (P). Representative of N = 5 donors. Q) LAE quantitation of a-tubulin area normalized to basement membrane length. N = 5 donors (represented by different colored dots). Paired t-test; ns = non-significant.

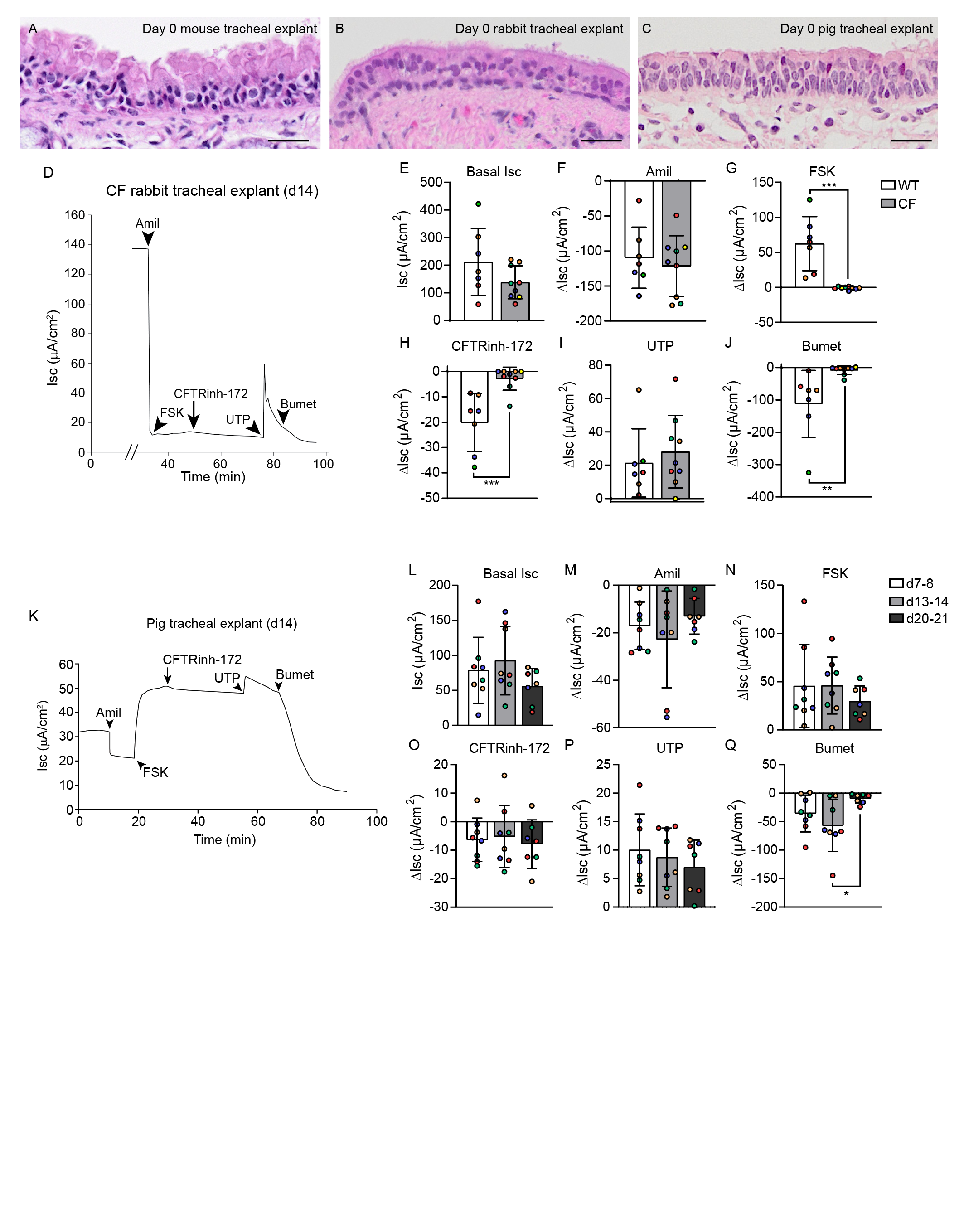

*Extended Data Fig 5.* A-C) H&E staining of day 0 mouse (A), rabbit (B), and pig (C) tracheal explants. Representative of N = 3, 2, and 3 animals, respectively. D-J) Electrophysiology of CF vs wildtype (WT) rabbit tracheal explants at d14. D) Representative Ussing tracing of a CF rabbit tracheal explant at d14. E) Basal Isc and ΔIsc in response to F) Amil, G) FSK, H) CFTRinh-172, I) UTP, and J) Bumet. N = 4-5 animals (represented by different colored dots); 1-2 replicates per animal. K-Q) Time course of pig tracheal explant electrophysiology at d7-8, d13-14, and d20-21. K) Representative Ussing tracing of a d14 pig tracheal explant. L) Basal Isc and ΔIsc in response to M) Amil, N) FSK, O) CFTRinh-172, P) UTP, and Q) Bumet. N = 4 animals (represented by different colored dots); 1-2 replicates per animal. One-way ANOVA with Tukey’s post-test; * = p<0.05.

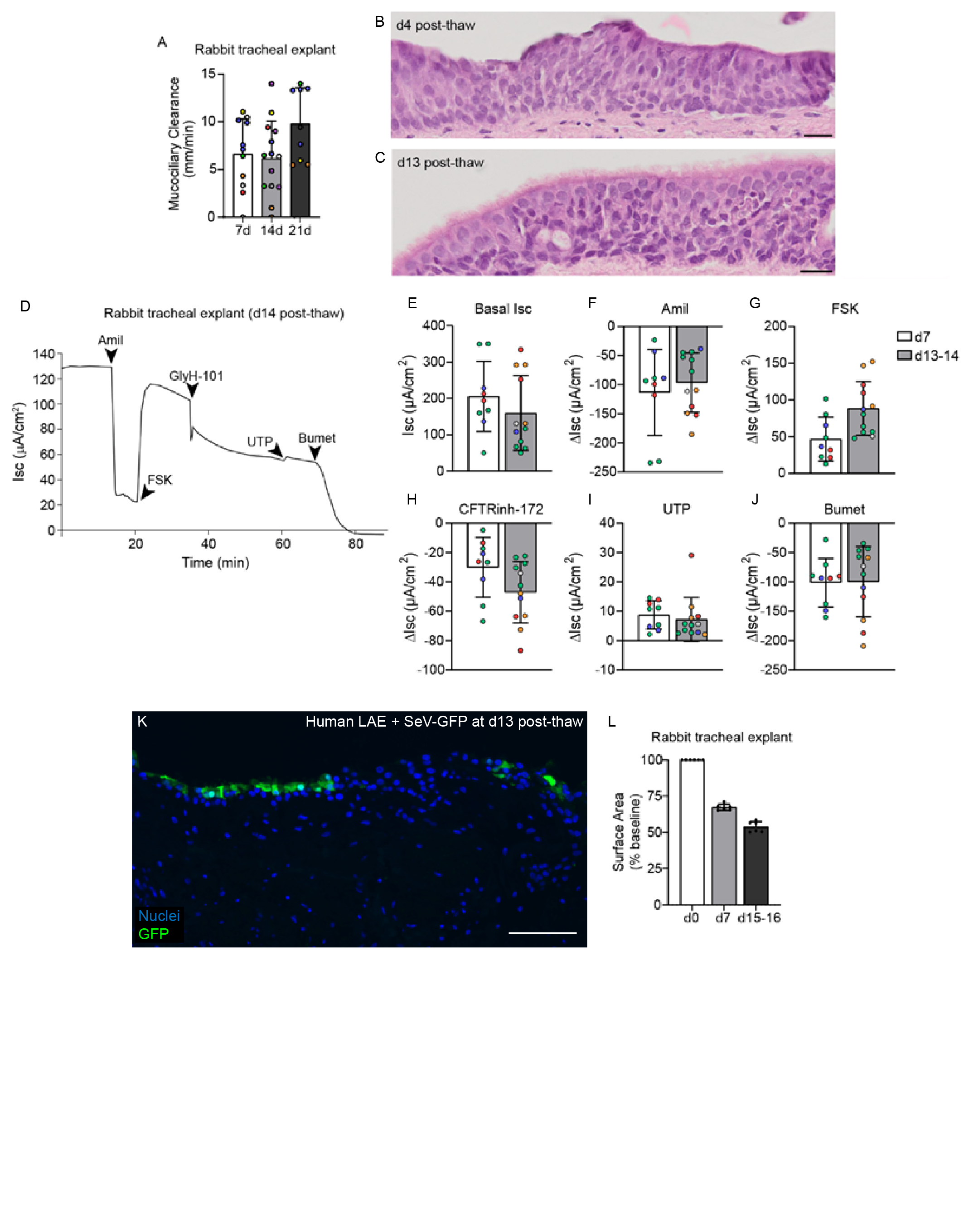

*Extended Data Fig 6. Rabbit tracheal explant mucociliary clearance and cryopreservation data.* A) Mucociliary clearance of fluorescently labeled beads across rabbit tracheal explants after 7, 14, and 21 days in culture. B-C) H&E histology of cryopreserved rabbit tracheal explants at 4 (B) and 13 (C) days post-thaw. Scale bars = 20 µm. D-J) Electrophysiology of cryopreserved rabbit tracheal explants at d7 and d13-14 post-thaw. D) Representative Ussing tracing of a cryopreserved rabbit tracheal explant at d14 post-thaw. E) Basal Isc and ΔIsc in response to F) Amil, G) FSK, H) CFTRinh-172, I) UTP, and J) Bumet. K) Immunofluorescence of a human LAE made from cryopreserved airway tissue at 3 days post infection (dpi) following SeV-GFP inoculation at d13 post-thaw. Scale bar = 100 µm. l) Surface area of rabbit tracheal explants over time as a percent of baseline.
